# Comparison of Brain Age Algorithms in Bipolar Disorder

**DOI:** 10.1101/2025.11.20.689600

**Authors:** Hui Xin Ng, Lisa Eyler

## Abstract

Advances in computational methods have accelerated the application of machine learning to analyze large complex biological data. By applying machine learning algorithms to neuroimaging data, researchers have estimated the “biological age of the brain” i.e., brain age, and used it as a composite metric for indexing brain health, as opposed to using individual features of the brain extracted from neuroimaging data. These machine learning algorithms/models, often known as “brain age” algorithms/models, may take supervised or unsupervised approaches and may utilize one or many imaging modalities during training. We applied 3 regression-based algorithm and 1 neural network-based algorithm trained on varying sample sizes of healthy comparison (HC) participants to estimate the brain age of 73 HC and 44 individuals with bipolar disorder (BD) in our neuroimaging study. Out of the four, 3 were pre-trained off-the-shelf algorithms and1 was developed and trained on multimodal neuroimaging data from a local cohort. The multimodal algorithm was trained on 51 age-matched HCs and tested on the remaining 22 HCs and 44 BDs. The brain predicted age difference (brain-PAD) score was calculated by subtracting the chronological age from the predicted age. Across four brain age prediction algorithms evaluated in HC, BrainageR and DenseNet demonstrated the highest predictive accuracy (r = 0.83; 0.89) and lowest mean absolute errors (MAE = 5.94; 7.26). However, PHOTON (r = 0.65, MAE = 7.71) showed greatest sensitivity to BD as demonstrated by our logistic regression model where the PHOTON brain-PAD was a significant predictor (beta = 0.064, p < 0.05) of BD. Analyses using ICC revealed that agreement levels varied, with PHOTON achieving the highest ICC with DenseNet (0.78) and BrainageR (0.73), which suggests they may pick up similar brain features as opposed to the multimodal algorithm (0.17- 0.43)

These results suggest that regularized linear models trained on large samples that explicitly exclude individuals with psychiatric diagnoses (i.e., PHOTON in this case) may be most sensitive to case-control differences despite having lower predictive accuracy. Our findings can serve as a starting point and quantitative reference for future efforts for researchers working with datasets that are similarly constrained by sample size but include unique combinations of imaging modalities.

## Introduction

### Aging and Modelling Aging in Bipolar Disorder

Bipolar disorder (BD) affects over 1% of the global population and is often misdiagnosed as major depressive disorder due to overlapping early symptoms. BD is linked to premature aging, with evidence from biomarkers such as oxidative stress (Andreazza et al., 2008; Berk et al., 2011), mitochondrial DNA copy number (Wang et al., 2018), and DNA methylation (Fries et al., 2017). Individuals with BD show elevated rates of age-related conditions and mortality (Hayes et al., 2015; Fiedorowicz et al., 2008), and a large study found their risk of dementia to be more than doubled (Nielsen et al., 2024). Longitudinal ENIGMA-BD data further demonstrate that greater manic episode burden predicts accelerated cortical thinning (Abé et al., 2022).

Given the shared features of mood disorders with aging in terms of brain structural alterations, regression-based, instead of classification-based, ML approaches have been applied to neuroimaging data to examine deviation from normative brain aging trajectories (Ballester et al., 2022; Lee et al., 2021; Seitz-Holland et al., 2024). This regression-based approach estimates the “biological age” at an individual level by comparing the individual’s brain scan against a normative population dataset of healthy individuals containing the same brain features.. Brain-redicted age difference (brain-PAD = predicted brain age - chronological age) is calculated by subtracting an individual’s chronological age from the predicted age. Brain-PAD is associated with age-related conditions and decline in physical functioning and mental health (Seitz-Holland et al., 2024; Wrigglesworth et al., 2021)

Smaller studies showed mixed findings on brain aging in BD: some found no difference compared to HC (Hajek et al., 2019, Nenadić et al., 2017, Shahab et al., 2019), while others found slight increases (Tønnesen et al., 2020) or lithium-associated reductions (Van Gestel et al., 2019). A meta-analysis demonstrated a modest overall brain-PAD increase in BD that ranges from +1.93 (fixed effects) to +2.12 years (random effects to account for variation in treatment effects across studies), despite moderate heterogeneity (Ballester et al. 2022). ENIGMA Consortium BD working group’s (under review; see Chapter 1) mega-analytic study on brain age in BD showed that individuals with BD have higher brain-PAD as age increases, although there was no main effect of BD status. Additionally, brain-PAD was the most pronounced among individuals taking antiepileptics or combined antiepileptic and antipsychotic medications. In another study of BD, lower brain-PAD was associated with lower global cognitive and verbal memory scores, which suggests neurodevelopmental delay in a relatively young cohort (Chakrabarty et al., 2022)

### Comparison of Machine Learning Pipelines

Baecker et al. (2021) provided a comprehensive overview of brain age prediction techniques and their clinical applications and highlighted the utility of pre-processed brain scans (e.g., voxel-wise features) in detecting neurodegeneration despite the increasing popularity of deep learning methods that utilize raw scans. Previous studies also have systematically classified deep learning architectures based on data type and modelling approaches (More et al., 2023, Tanveer et al. 2023), evaluated how different brain-age prediction workflows varying in feature representations, training dataset and ML algorithms perform, i.e., within-dataset accuracy, cross-dataset generalization, test-retest reliability, and longitudinal consistency. (Bacas et al., 2023; Dörfel et al., 2023; Soumya, Kumari & Sundarrajan, 2024; Yu et al., 2024) and compared traditional ML approaches to emerging techniques (Soumya Kumari & Sundarrajan, 2024).

Assessing out-of-sample generalizability is challenging because while brain age algorithms trained on large diverse datasets have shown high cross validation accuracy, their performance on new, independent samples with different demographics and imaging characteristics remains unclear, because good performance during cross validation overestimates generalizability (de Lange et al, 2022; Poldrack et al., 2020). Bashyam et al. (2020) showed that overfitted algorithms (e.g. algorithms with good performance in training and cross validation) are not the most sensitive in identifying pathologies; moderately fit algorithms are better at detecting group differences at large effect sizes between control and patient groups across brain disorders.

### Brain age algorithms have demonstrated utility for diagnosis and prognosis and have been linked to deviations in cognitive performance and disease progression

1 data from a small local sample could yield meaningful insights and demonstrate added value beyond existing pre-trained algorithms. Specifically, we compared the following algorithms as applied to a sample of people with and without BD: i) PHOTON, developed by the ENIGMA-MDD group (L. K. M. Han, Dinga, et al., 2021) ii) brainageR developed by MANIFOLD lab (https://github.com/james-cole/brainageR), iii) DenseNet developed by MANIFOLD lab (Wood et al, 2022) a new Multimodal algorithm developed locally.

**Table 2.1.** Characteristic of Brain Age Algorithms

| Characteristic | Description |
| --- | --- |
| Feature Representation | The form and structure of the brain imaging data used as input—this can include region-based metrics (e.g., cortical thickness), voxel-wise maps, or multimodal imaging features. Coarse features (e.g., cortical thickness) may miss fine-grained pathology; voxel-wise or multimodal features may capture richer variation. |
| Training Dataset Size | The number of healthy individuals used to train the model. Larger datasets generally improve generalizability and prediction accuracy. |
| Age Range | Determines the age-related variance the model can learn; too narrow a range limits generalizability across the lifespan. A narrow age range can cause bias (e.g., poor predictions at age extremes) |
| Performance on Test Dataset | The model’s accuracy on a held-out test sample, often measured by Mean Absolute Error (MAE) and Pearson correlation (r) with chronological age. high accuracy doesn't guarantee clinical utility—some high-performing algorithms may miss disease-related deviations (underfit pathology) |
| Training Dataset Modality | The type of MRI imaging modalities used to extract features—e.g., T1-weighted, T2-weighted, FLAIR, DTI, ASL, and fMRI data. |

Algorithms were evaluated based on mean absolute error (MAE) and Pearson’s correlation (r) between predicted and chronological age. In addition, model agreement was assessed using intraclass correlation coefficients (ICC).

The Multimodal algorithm, despite being trained on a smaller sample, was expected to be sensitive to BD given its richness in features across imaging modalities. In contrast, we expected the brainageR and DenseNet algorithms, which were trained on larger and more homogeneous datasets, to yield higher prediction accuracy in healthy controls. We also hypothesized that ICC, which measures the level of agreement between algorithms, would be highest between DenseNet and brainageR, as both were trained on large, T1-weighted datasets with high granularity and image resolution, and lowest between the Multimodal algorithm and the others, due to differences in input features, preprocessing, and training sample characteristics.

## Methods

### Multimodal Algorithm Dataset Sample

Participants were 44 individuals with BD I (stable mood, first episode age 13–30) and healthy controls, screened to exclude psychiatric, neurological, or substance use disorders. All completed the D-KEFS cognitive battery and were eligible for MRI. Cognitive testing coincided with scanning for some individuals, while blood draws occurred separately (mean delay = 82 days). while blood draws were performed on a different day, with an average delay of 82 days. All procedures were approved by the Institutional Review Board of the San Diego Veterans Affairs Healthcare System. See Dev et al. 2017 for image acquisition and data processing protocols.

### Machine Learning Model Types

**Table 2.2.**
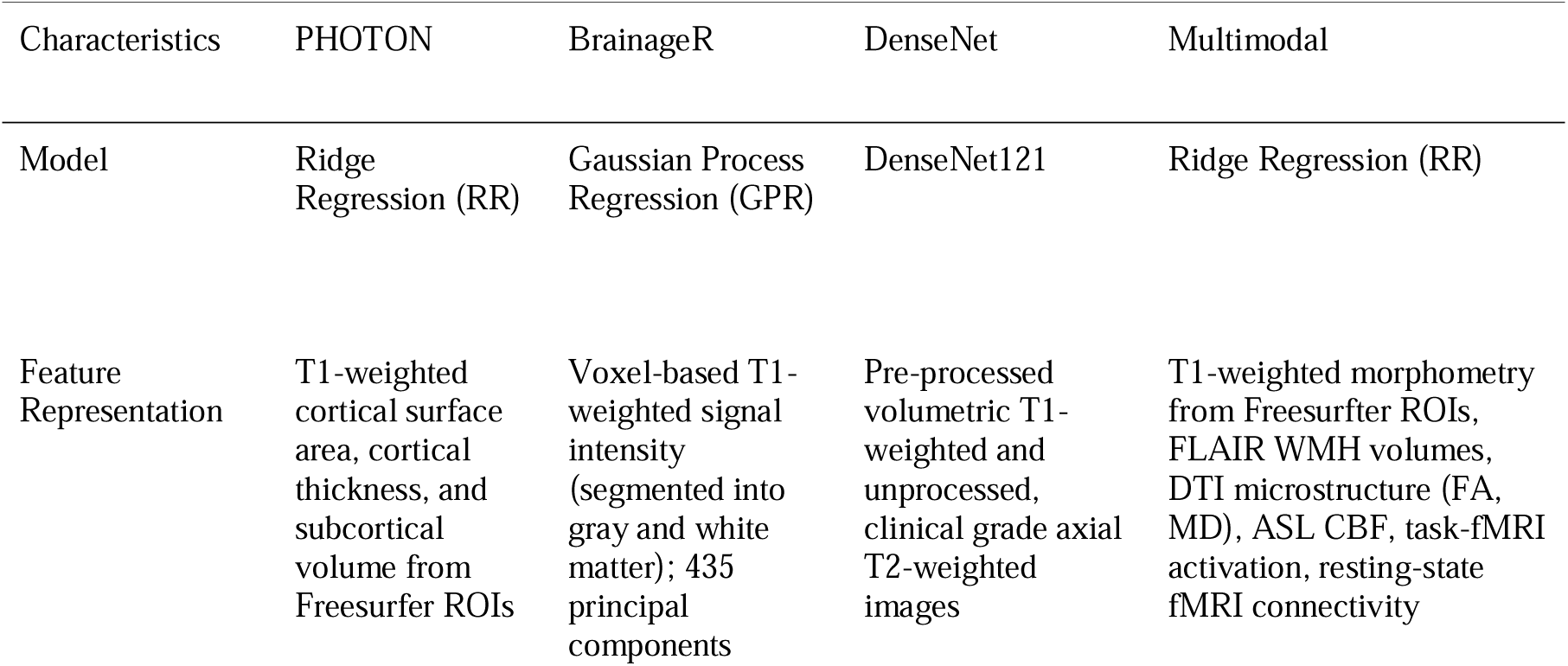
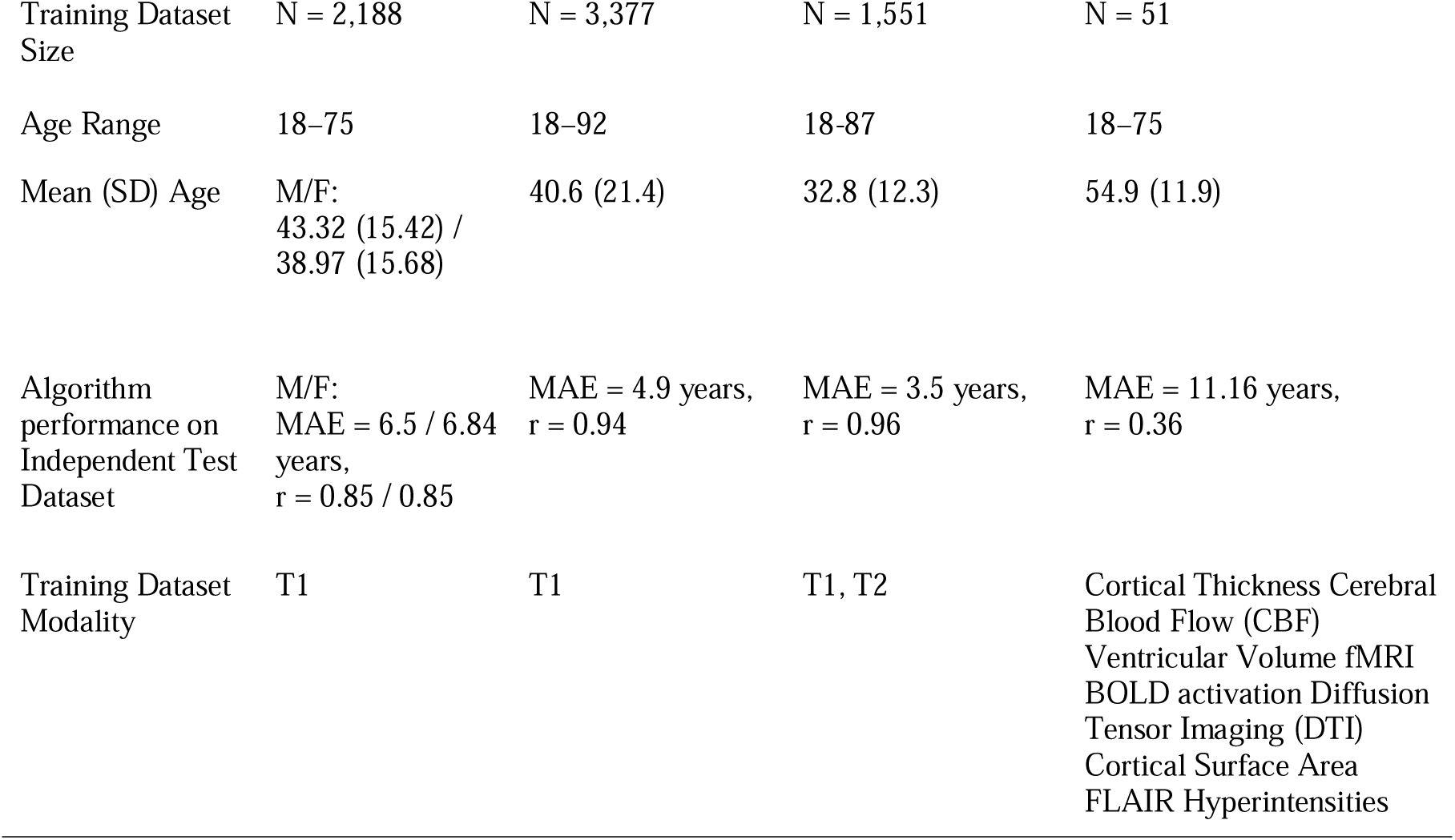
Comparison of four brain age algorithms, detailing model type, feature representation, training sample size and age range, independent test performance (MAE and Pearson’s r) and imaging modality.

We examined four machine learning algorithms to obtain predicted brain age and brain-PAD values: i) PHOTON, ii) brainageR, iii) DenseNet, and iv) Multimodal algorithm. PHOTON (https://www.photon-ai.com/enigma_brainage) and Multimodal algorithm were trained on FreeSurfer-parcellated cortical and subcortical features, brainageR (https://zenodo.org/record/3476365#.ZFBPvHbMKUk) was trained on voxel-based T1-weighted scans. Wood et al (2022) provided a version of the DenseNet algorithm that was trained pre-processed volumetric T1- weighted and unprocessed, clinical grade axial T2-weighted images using a 3D ensemble convolutional neural network. If only T1-weighted scans are available as input, the ensemble model defaults to T1-based prediction alone, enabling model utility in single-modality datasets.. PHOTON, brainageR, and DenseNet MIDI are pre-trained algorithms, while Multimodal ridge was developed locally using our dataset consisting of multimodal imaging features. suited to capture complex patterns in brain imaging data, compared to linear algorithms.

### Details about pre-trained algorithms

PHOTON ridge regression model, trained on FreeSurfer-derived T1 features from 2,188 healthy controls, has demonstrated robust generalizability and has been widely applied across ENIGMA working groups to compute brain-PAD (L. K. M. Han, Dinga, et al., 2021; Clausen et al., 2022; Constantinides et al., 2022) The brainageR Gaussian Process Regression model, trained on 3,377 T1 scans from seven public datasets, achieved high predictive accuracy with no age bias in external validation and has been linked to mortality risk (Cole et al., 2018;).DenseNet-121, a 3D CNN trained on ∼23,000 hospital scans was able to predict brain age with high accuracy, generalized across sites/scanners, and identified clinically relevant age-excessive atrophy. See Table 2.2 for more details.

### Development of Multimodal algorithm using our local dataset

For our local algorithm, we selected ridge regression (RR) given its simplicity, interpretability, and strong generalizability in smaller neuroimaging samples. RR is a linear model with L2 regularization that shrinks coefficients to mitigate overfitting, particularly effective in high-dimensional settings with multicollinearity among features. The optimal regularization strength was selected via repeated five-fold cross-validation (10 repetitions), and model performance was evaluated using MAE and Pearson’s correlation. Further methodological details are provided in the Supplementary Materials (Supplementary Figures 2.1–2.2).

Age bias correction: Except for brainageR, all algorithms showed significant correlations between chronological age and brain-PAD (Supplementary Figures 2.6–2.8), reflecting the regression-to-mean effect whereby younger ages are overestimated and older ages underestimated (Cole, 2020; de Lange & Cole, 2020; Le et al., 2018; Smith et al., 2019). For brainageR, no significant correlation was observed (r = –0.05, p = 0.56), nor did age interact with sex or diagnosis, so age was not included as a covariate in the final model.

Algorithm performance comparison: Analyses were conducted in R and Python (scikit-learn), evaluating algorithm performance in healthy controls using R², Pearson’s r, and MAE, with agreement assessed via pairwise ICC(3,1).

Statistical analysis: To test age and sex moderation of BD–HC brain-PAD differences, we used stepwise models including all interactions (Dx × Age × Sex). Age and sex were included as covariates given known effects on brain structure (Mitchell et al., 2016; Ziemka-Nalecz et al., 2023; Almeida et al., 2022; Blanken et al., 2024; Xu et al., 2021) and to adjust for age-related bias unless brain-PAD was uncorrelated with chronological age.

**Table 2.3.**
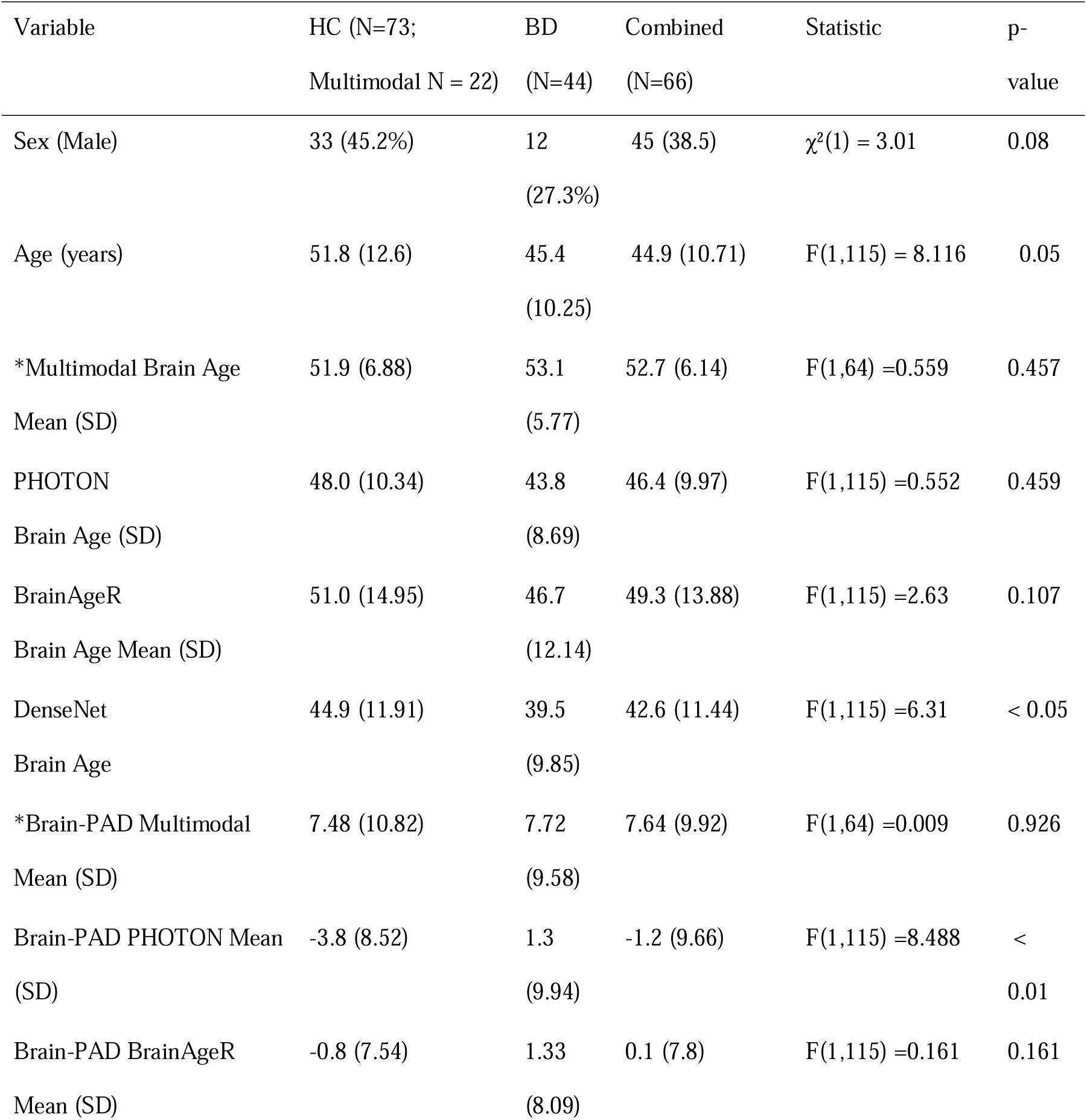
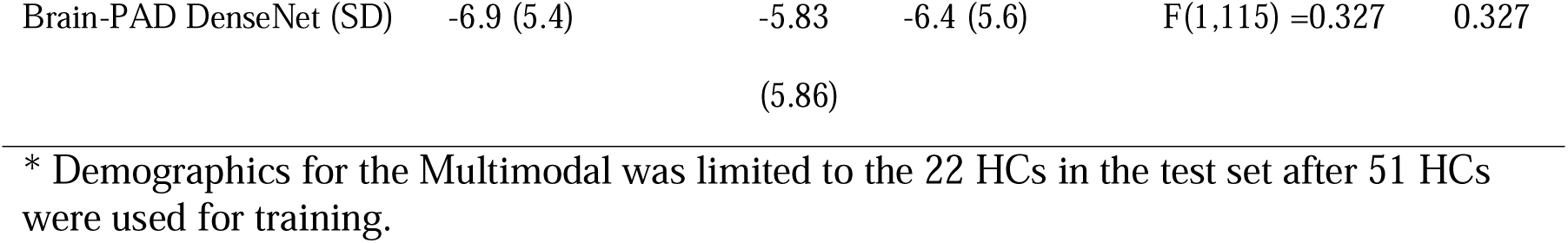
Sample demographic and clinical characteristics

## Results

### Algorithm performance

Among HC, the DenseNet model achieved the best overall performance, with a mean absolute error (MAE) of 7.26 years, an R² of 0.81, and a Pearson’s correlation coefficient (r) of 0.90. BrainageR also demonstrated strong performance, achieving the lowest MAE of 5.94 years, an R² of 0.74, and r = 0.86. PHOTON showed moderate performance with an MAE of 7.71 years, an R² of 0.55, and r = 0.74. In contrast, the Multimodal algorithm exhibited the weakest performance, with the highest MAE (11.16 years) and the lowest correlation with chronological age (r = 0.38, R² = 0.42). See supplementary Figure 2.5-2.7 for visualization of the correlations.

**Table 2.4.** MAE, Pearson’s correlation and R^2^ across PHOTON, DenseNet, BrainageR and Multimodal algorithms

|  | PHOTON<br>(HC = 73) |  |  | BrainageR<br>(HC = 73) |  |  | DenseNet<br>(HC = 73) |  |  | Multimodal |  |  |
| --- | --- | --- | --- | --- | --- | --- | --- | --- | --- | --- | --- | --- |
| Metric | MAE | $R^2$ | r | MAE | $R^2$ | r | MAE | $R^2$ | r | MAE | $R^2$ | r |
|  | 7.71 | 0.55 | 0.74 | 5.94 | 0.74 | 0.86 | 7.26 | 0.81 | 0.90 | 11.16 | 0.42 | 0.38 |

In terms of agreement between algorithms, when HC and BD groups are combined, the strongest agreement was observed between DenseNet and PHOTON (ICC = 0.78) and between PHOTON and BrainageR (ICC = 0.73). Multimodal demonstrated the weakest agreement with other algorithms, particularly with DenseNet (ICC = 0.17), suggesting greater divergence in predicted brain ages (See supplementary Table 2.8 for 95% CI of the ICC). The ICC demonstrates consistency and absolute agreement between the predicted ages, capturing not only how well they co-vary but also any systematic differences in level or scale. We included the Pearson’s r correlations across the predicted ages across the algorithms in Supplementary material Figure 3.2. Overall, ICC agreement between algorithms was higher in HC compared to individuals with BD. In HC, PHOTON and DenseNet showed the strongest agreement (ICC = 0.80), followed by PHOTON and BrainageR (ICC = 0.76). In BD, agreement was lower across all comparisons, with the highest ICC again observed between DenseNet and PHOTON (ICC = 0.70). Multimodal consistently demonstrated the weakest agreement with other algorithms, particularly with DenseNet (ICC = 0.19 in HC and 0.16 in BD), suggesting that its predicted brain ages diverged substantially from those of other algorithms.

**Figure 2.1.**
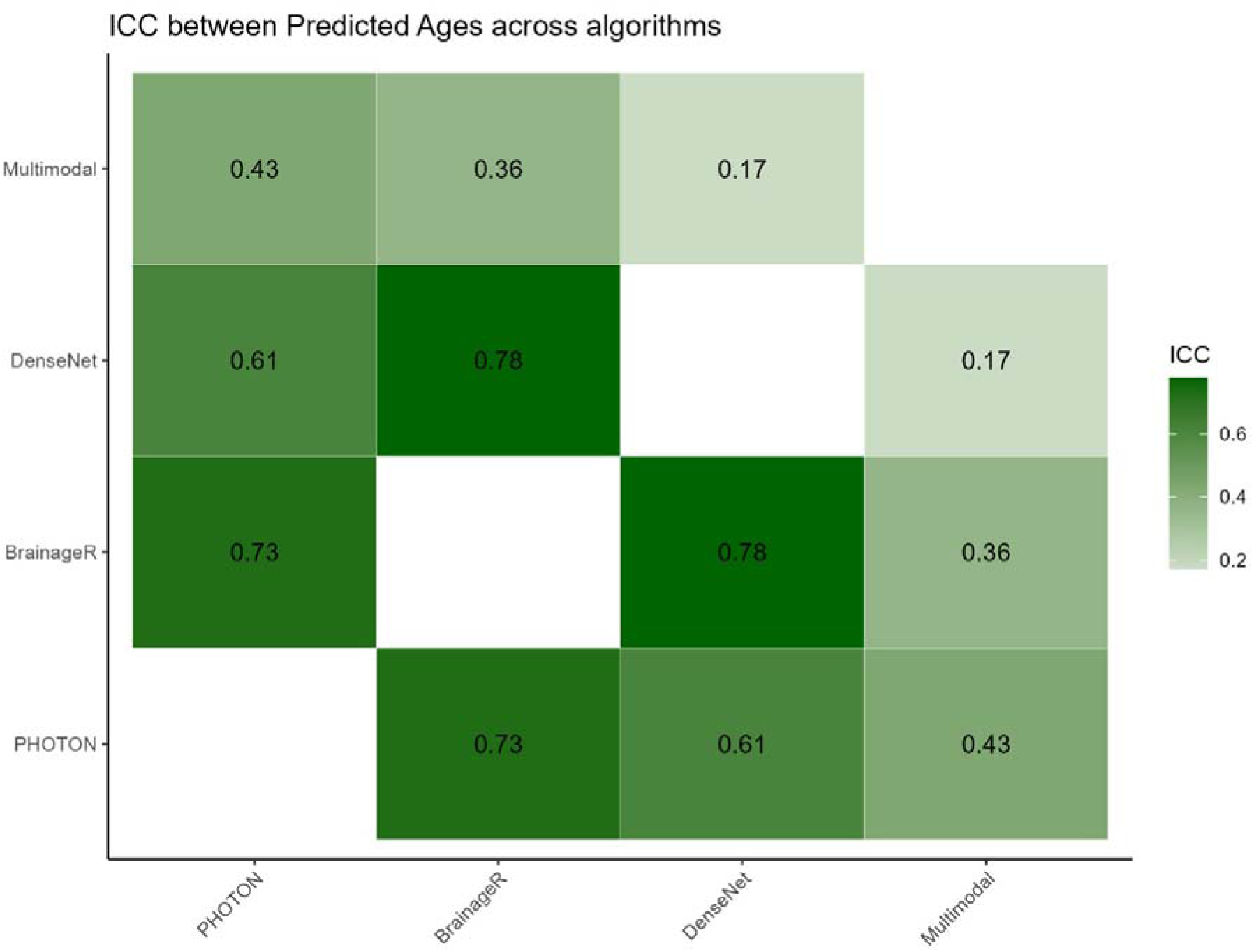
Pairwise intraclass correlation coefficients (ICCs) between predicted brain ages generated by four algorithms: PHOTON, BrainageR, DenseNet, Multimodal

### Group differences and moderation analysis

**Table 2.5.**
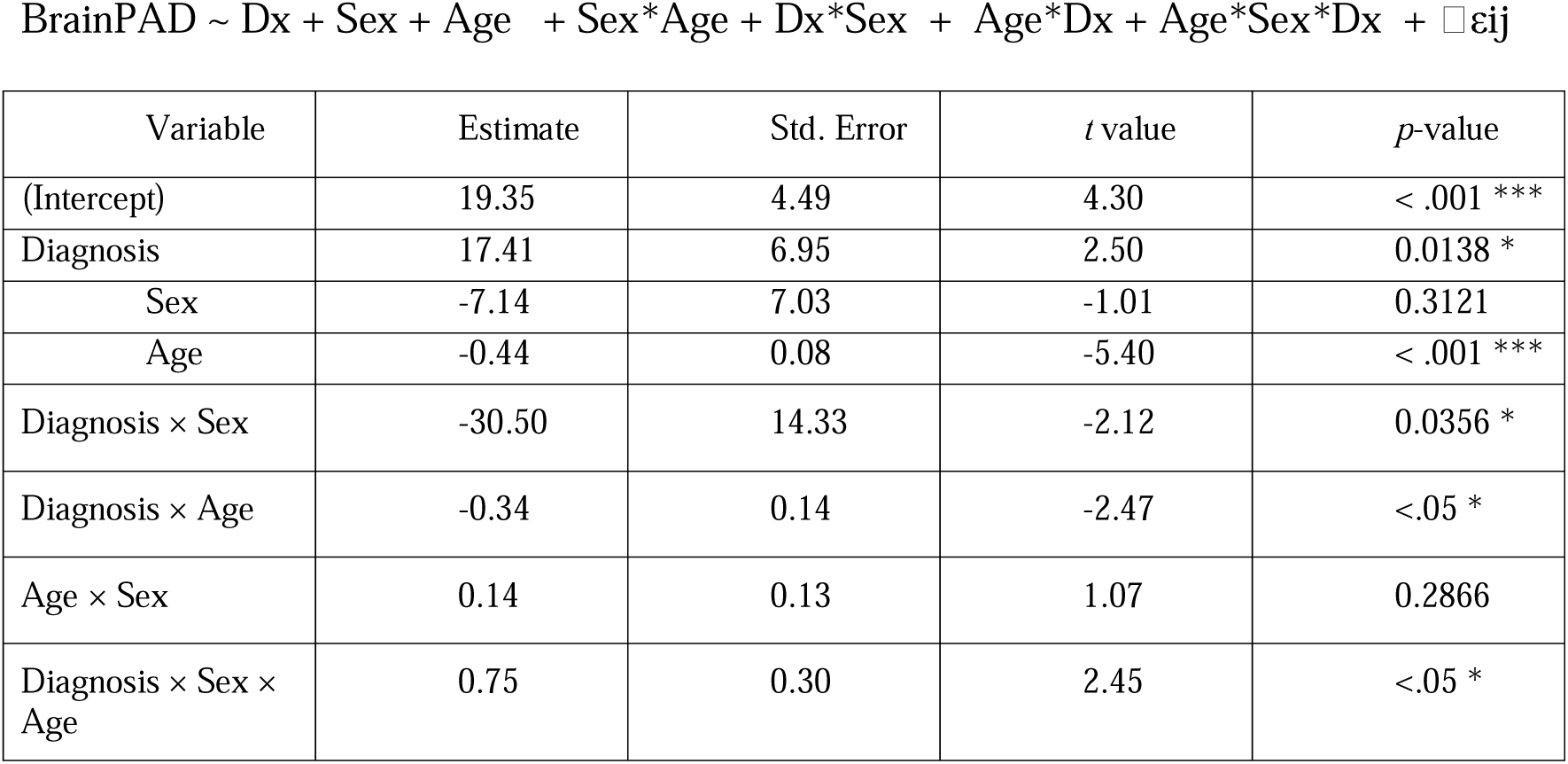
Selected linear regression model fit to brain-PAD produced from PHOTON:

**Table 2.5.** Selected linear regression model fit to brain-PAD produced from BrainageR:

| Variable | Estimate | Std. Error | *t* value | *p*-value |
| --- | --- | --- | --- | --- |
| Intercept | -1.867 | 1.1228 | -1.6633 | 0.098994 |
| Diagnosis | 2.525 | 1.4915 | 1.6930 | 0.093172 |
| Sex | 2.461 | 1.4851 | 1.6577 | 0.100123 |

**Table 2.6.** Selected linear regression model fit to brain-PAD produced from DenseNet:

| Variable | Estimate | Std. Error | *t* value | *p*-value |
| --- | --- | --- | --- | --- |
| Intercept | 0.3732 | 2.2947 | 0.1626 | 0.8711 |
| Diagnosis | 0.3835 | 1.0462 | 0.3666 | 0.7146 |
| Sex | 1.9031 | 1.0070 | 1.8899 | 0.0613 |
| Age | -0.1567 | 0.0412 | -3.8005 | 0.0002 |

**Table 2.7.**
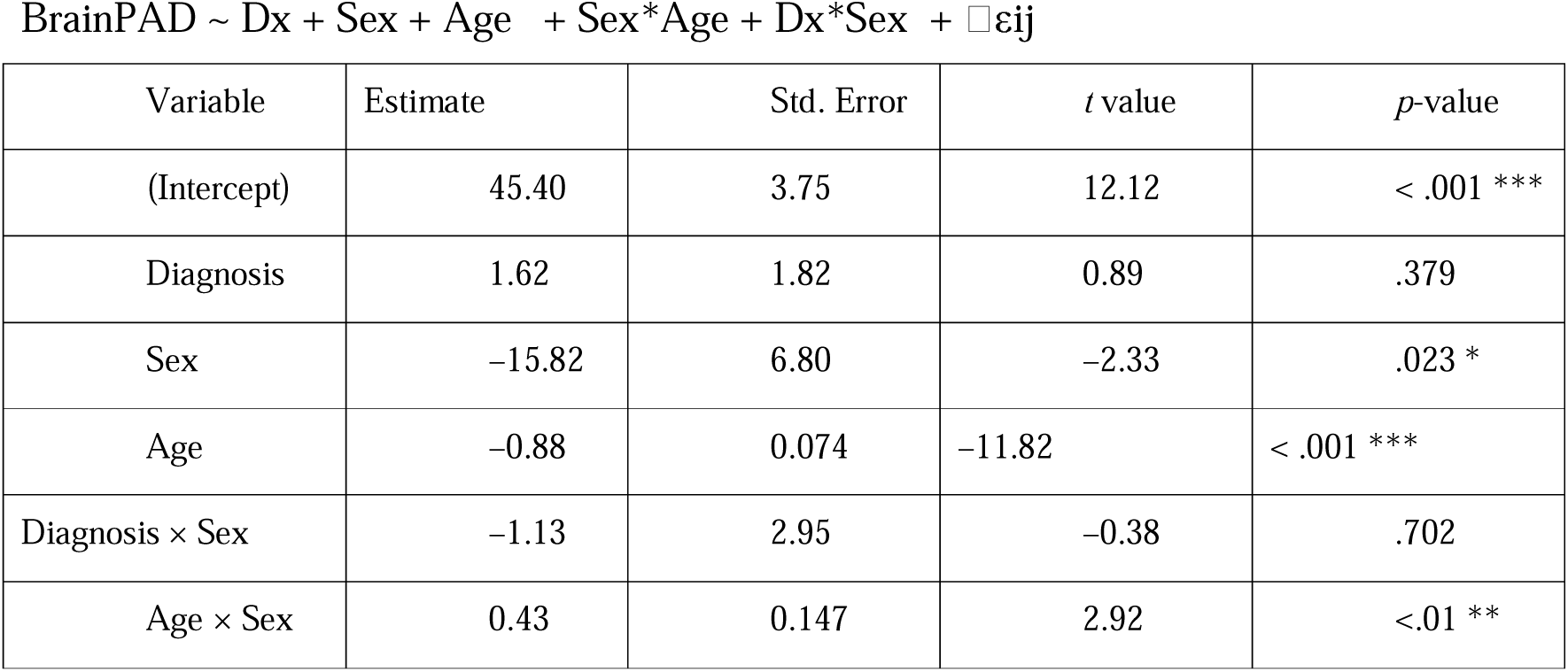
Selected linear regression model fit to brain-PAD produced from Multimodal:

The PHOTON algorithm uniquely detected group differences, with a significant diagnosis × sex × age interaction (β = 0.75, t = 2.45, p = .014), showing brain-PAD in BD men increased 0.75 years faster per chronological year than in BD women. PHOTON also showed main effects of diagnosis (β = 17.41), age (β = –0.44), diagnosis × sex (β = –30.50), and diagnosis × age (β = –0.34).For BrainageR and DenseNet, best-fit algorithms included only main effects, with no significant BD effect (BrainageR: β = 2.52, t = 1.69, p = .09; DenseNet: β = 0.38, t = 0.37, p = .72).The Multimodal algorithm retained main effects plus two interactions. Significant effects were age (β = –0.88, t = –11.82, p < .001), sex (β = –15.82, t = –2.33, p = .023), and sex × age (β = 0.43, t = 2.92, p = .005). Diagnosis × sex was nonsignificant (p = .702) but retained for better fit (χ²(1) = 247.36, p = .0035).

## Discussion

In the present study, we analyzed the predictive power and reliability of brain age algorithms that are publicly available, trained on large sample sizes and span across diverse algorithm types: (PHOTON, BrainageR, DenseNet), in addition to our locally trained algorithm (Multimodal). Although the multimodal algorithm had several limitations, developing and testing it on data from the same cohort and scanner is a potentially valuable approach that enables researchers with small sample sizes and a specific set imaging modalities that are typical of local neuroimaging datasets to assess whether they can derive meaningful brain-PAD estimates by developing their own algorithms.

Standardization of benchmarks for model/algorithm development and evaluation remains an open question in the field of research on brain age algorithms. While many benchmark analyses have been attempted to compare the accuracy and reliability of various algorithms (Dular & Špiclin, 2024, Dufumier et al., 2022,Dörfel et al., 2023), the field still needs to address the challenge of validating brain-PAD and predicted age as biomarkers, as many have pointed out that most accurate algorithms, as most frequently measured through MAE and correlation between predicted and chronological ages, may not meaningfully model information needed to distinguish between HCs and individuals with diagnoses (Jirsaraie et al., 2022, 2023).

### Algorithm performance and agreement

Our results indicate that dataset with richer feature representations (e.g. whole-brain voxels or CNN-extracted features) and non-linear modeling capacity are critical for predictive accuracy. However, the superior performance of DenseNet is at the cost of transparency and interpretability. Algorithms using scans that are pre-processed (Freesurfer parcellated brain regions or PCAs) are more interpretable despite lower accuracy. Beyond the trade-off between predictive accuracy and interpretability in brain age prediction, prior work has also shown algorithms that predict chronological age less accurately, such as deep learning algorithms that are loosely fitted (Bashyam et al., 2020) and linear algorithms that incorporate regularization, can pick up case-control differences with greater sensitivity (Schulz et al, 2024).

Our ICC analysis reveals a wide range of agreement levels across the algorithms, with PHOTON achieving the highest ICC with DenseNet and BrainageR, which indicates there is an overlap in their ranking and the exact values of the predicted ages. All three algorithms rely on structural imaging features, albeit processed differently. It is unsurprising that ICC between DenseNet and Multimodal was the lowest (0.17), since the difference in modality and input representation diverges the most. This divergence reflects the contrast in input representation and algorithm type between the Multimodal and DenseNet algorithms; the prior combines multiple imaging modalities and is a regression model, whereas the latter is a convolutional neural network on T1-weighted MRI. Each model may emphasize distinct aspects of brain structure or pathology, in addition to capturing error and natural variability differently. In other words, the algorithms with high mutual ICC measure more similar brain aging patterns, while the low-ICC pair suggests that those two algorithms are not directly comparable and may be capturing different facets of aging or subject to different biases.

The PHOTON algorithm was the only algorithm that showed BD–HC differences, as demonstrated by significant age × sex interactions. PHOTON has also generalized well in ENIGMA MDD mega-analyses (>3400 patients, >2800 controls), where MDD was associated with +1 year brain-PAD (Han et al., 2021; 2022). Training composition influences sensitivity: DenseNet was trained on radiologically “normal” scans without excluding mood disorders, and brainageR included datasets (e.g., IXI, NKI) that may have contained individuals with mood disorders, potentially embedding disorder-related features into the normative baseline. In contrast, models with stronger regularization show greater sensitivity to pathology; Schulz et al. (2024) reported ridge regression detected the largest brain-PAD effects in UK Biobank data (>46,000), with effect size diminishing as model complexity increased. This suggests simpler, interpretable models may yield more robust biomarkers of psychiatry-related pathology (Han et al., 2021; He et al., 2020; Schulz et al., 2024).

#### Future Directions and Limitations

Best practices for evaluating brain-age models remain unsettled, particularly regarding whether they capture true normative aging (Cole et al., 2019; Nguyen et al., 2024; Vidal-Pineiro et al., 2021; Wrigglesworth et al., 2022). Most models, including ours, are trained on cross-sectional data, limiting their ability to reflect individual trajectories shaped by genetics and environment. While cross-sectional models can reveal group-level case–control differences, longitudinal assessments are essential for quantifying individual rates of change and prediction error. Normative modeling approaches instead characterize aging trajectories as a function of covariates (e.g., age, sex) and define individual deviations in brain-PAD. Shifting toward these individualized trajectories may identify BD subgroups with accelerated aging, even when mean group differences are absent. Importantly, model utility also depends on phenotype: large deviations in brain-PAD are seen in illnesses with progressive atrophy (e.g., Alzheimer’s, schizophrenia; Jirsaraie et al., 2023; Dias et al., 2025). Our findings add to validation of brain-PAD as a potential biomarker in BD.

Our study is constrained by a small sample (HC = 73; BD = 44), limiting scalability, generalizability, and statistical power. The multimodal algorithm was tested in only 22 HCs, and apparent differences (e.g., PHOTON’s sensitivity to BD) may reflect sample idiosyncrasies. Although we lacked longitudinal data, prior work shows high reliability for brainageR but poorer stability for PHOTON (Dörfel et al., 2023; Hanson et al., 2024). We also did not examine clinical moderators such as medication or cognition, despite evidence linking elevated brain-PAD to progression and decline (Beheshti, 2025). Future studies should validate subgroup-specific trajectories, assess moderators, and clarify whether brain-PAD reflects pathological rather than normative aging. Despite these caveats, our results suggest that deep learning and linear models capture overlapping structural patterns, whereas multimodal models diverge, offering guidance for local model development.

