## Supplemental Material for "Comparison of Brain Age Algorithms in Bipolar Disorder"

Supplemental Table 2.1. Demographics and brain age-related metrics for smaller HC test set (N=22) held out during Multimodal algorithm development

| Variable | HC (N=22) | BD (N=44) | Combined (N=66) | Statistic | p-value |
| --- | --- | --- | --- | --- | --- |
| Sex (Male) | 10 (45.5%) | 12 (27.3%) | 22 (33.3%) | χ²(1) = 1.44 | 0.23 |
| Age (years) | 44.4 (11.14) | 45.4 (10.25) | 44.9 (10.71) | F(1,64) =0.32 | 0.57 |
| PHOTON Brain Age (SD) | 43.8 (8.69) | 43.8 (8.69) | 45.7 (8.44) | F(1,64) =1.639 | 0.205 |
| BrainAgeR Mean (SD) | 44.1 (11.72) | 46.7 (12.14) | 45.8 (11.97) | F(1,64) =0.684 | 0.411 |
| DenseNet Brain Age  Mean (SD) | 38.3 (10.92) | 39.5 (9.85) | 39.1 (10.15) | F(1,64) =0.210 | 0.648 |
| Multimodal Brain Age  Mean (SD) | 51.9 (6.88) | 53.1 (5.77) | 52.7 (6.14) | F(1,64) =0.559 | 0.457 |
| Brain-PAD PHOTON Mean (SD) | -0.55 (9.2) | 1.3 (9.94) | 0.68 (9.67) | F(1,64) =0.533 | 0.468 |
| Brain-PAD BrainAgeR Mean (SD) | -0.31 (7.57) | 1.33 (8.09) | 0.78 (7.9) | F(1,64) =0.625 | 0.432 |
| Brain-PAD DenseNet MIDI (SD) | -6.1 (6.95) | -5.83 (5.86) | -5.92 (6.19) | F(1,64) =0.026 | 0.871 |
| Brain-PAD Multimodal Mean (SD) | 7.48 (10.82) | 7.72 (9.58) | 7.64 (9.92) | F(1,64) =0.009 | 0.926 |

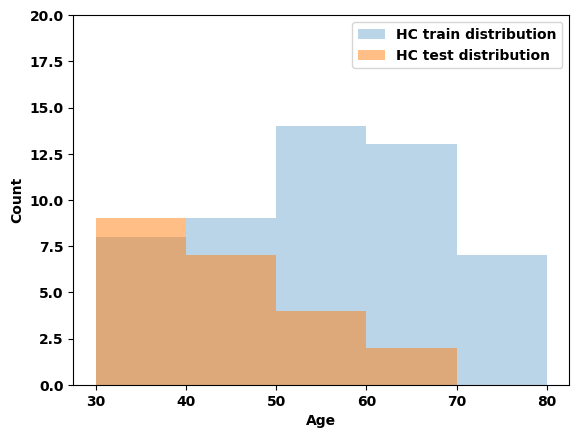

Supplemental Figure 2.1 Distribution of Age Training and Test Sets of HC

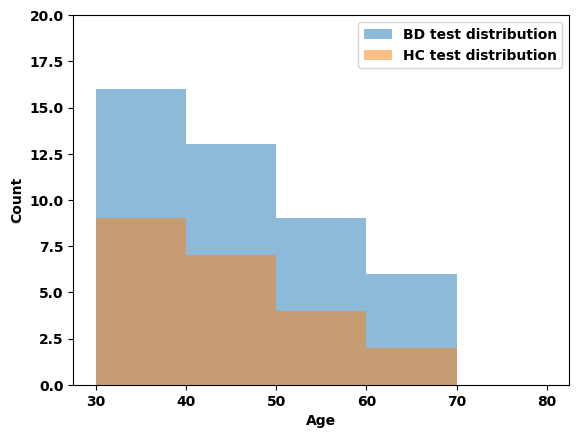

Supplemental Figure 2.2 Distribution of Age Test Set of HC and BD

*Characteristics of training and test sets*

We applied a regression-based machine learning framework to predict chronological age from multimodal neuroimaging data using ridge regression (RR) and support vector regression (SVR), selected for their robustness in small-sample, high-dimensional settings. Features (See Supplemental Table 2.1) from seven imaging modalities (DTI, CBF, cortical thickness, cortical surface area, ventricular volume, fMRI BOLD activation, FLAIR hyperintensities) were standardized using MinMax or Z-score normalization. The model was trained on a subset of 51 HCs and tested on 44 BD and 22 HC individuals. Supplemental Figure 2.1 shows the distribution of HC training dataset compared to the distribution of the HC test dataset when the model is trained on the subset of the HC group; Supplemental Figure 2.2 shows the distribution of the HC test dataset compared to the BD test dataset. This illustrates that the training distribution covers the entire age range, while fulfilling the requirement that age distribution A chi-squared test was conducted to compare the observed frequencies of participants across five age groups (30–39, 40–49, 50–59, 60–69, and 70–79) within the healthy control (HC) training set against expected frequencies. The results indicated no significant deviation from the expected distribution, χ²(𝑑𝑓) = 3.80, p = 0.43. This suggests that age group representation in the HC training set is approximately uniform and not biased toward any particular age group.

*Training and validation*

Models were trained on healthy controls using 5-fold cross-validation repeated 10 times. Hyperparameters were tuned via internal cross-validation: RR used a grid of alpha values, while SVR parameters (C, epsilon, kernel) were optimized via grid search. To evaluate the contribution of each modality, we implemented a leave-one-modality-out ablation procedure, retraining the model each time and assessing algorithm performance with each iteration. Algorithm performance was assessed using mean absolute error (MAE) and Pearson correlation between predicted and chronological age. The MAEs are presented in Table X for both RR and SVR with each leave-one-modality-out iteration.

*Performance of leave-one-modality-out models*

Both models were evaluated for brain age prediction using cross-validation on the training set and an independent test set for validation. The ridge regression model achieved a cross-validation MAE of 8.21 years and a test set MAE of 11.17 years.  Similarly, the support vector regression (SVR) model yielded a cross-validation MAE of 8.26 years and a test set MAE of 11.73 years. While both models showed comparable performance during cross-validation, the higher MAE on the test set suggests a moderate decline in performance when generalizing to new, unseen data. RR was chosen given its lower CV MAE and overall MAE. We refer to this model as Multimodal throughout the paper.

Supplemental Table 2.2 MAE for Ridge Regression and Support Vector Regression models in a leave-one-modality-out analysis across different brain imaging modalities.

| Leave-one-modality-out | Mean Absolute Error (MAE) | |
| --- | --- | --- |
| Ablated Feature Set | Ridge Regression | Support Vector Regression |
| Cortical Thickness | 9.74 | 9.65 |
| Cerebral Blood Flow (CBF) | 9.95 | 10.35 |
| Ventricular Volume | 10.01 | 10.35 |
| fMRI BOLD activation | 10.13 | 10.27 |
| Diffusion Tensor Imaging (DTI) | 10.51 | 11.24 |
| Cortical Surface Area | 10.20 | 10.66 |
| FLAIR Hyperintensities | 10.58 | 10.66 |

Supplemental Table 2.3. Algorithm performance Metrics (MAE, R2, r) across four algorithms when applied to full sample of HC (N = 73) and subset of HC (N=22)

|  | PHOTON | | | BrainageR | | | DenseNet | | | Multimodal | | |
| --- | --- | --- | --- | --- | --- | --- | --- | --- | --- | --- | --- | --- |
| HC N | MAE | R^2^ | r | MAE | R^2^ | r | MAE | R^2^ | r | MAE | R^2^ | r |
| (HC = 73) | 7.71 | 0.55 | 0.74 | - | 0.74 | 0.86 | 7.26 | 0.81 | 0.90 | - | - | - |
| (HC = 22) | 7.48 | 0.35 | 0.59 | 11.16 | 0.61 | 0.78 | 6.72 | 0.79 | 0.89 | 11.16 | 0.42 | 0.38 |

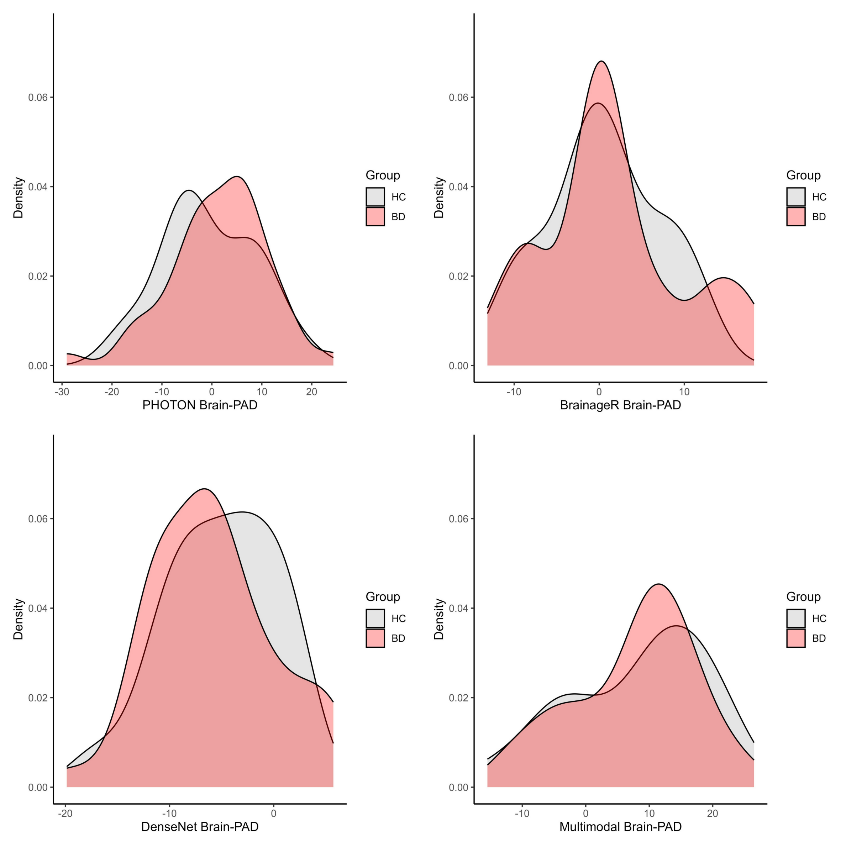

Supplemental Figure 2.1. Brain-PAD Distribution of Brain-PAD produced from PHOTON, BrainageR, DenseNet, and Multimodal algorithms

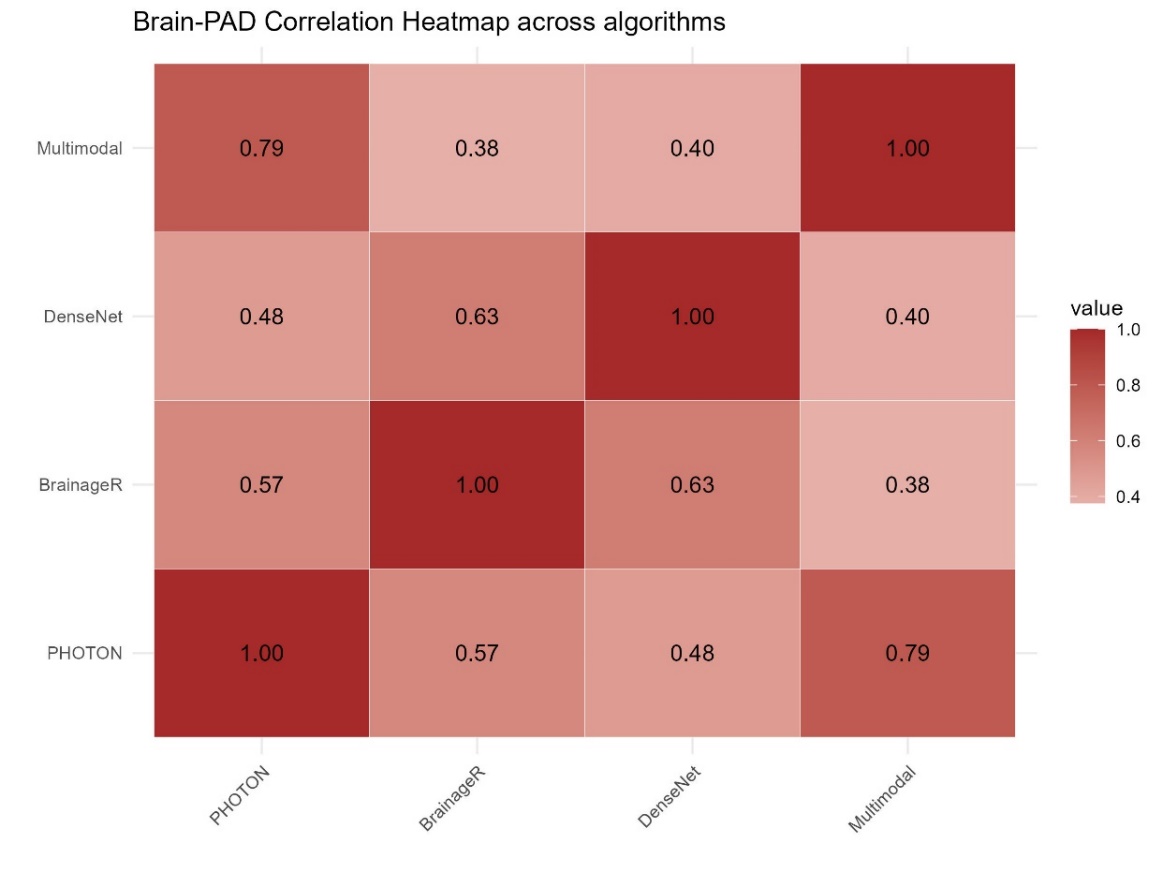

Supplemental Figure 2.3. Pearson’s correlation between predicted ages across algorithms

*Agreement among algorithms as assessed by correlation patterns*

Across both HC and BD groups, correlation patterns varied depending on whether raw predicted ages or brain-PAD values were analyzed. In general, predicted ages showed moderate to high correlations across models (r = 0.46–0.89), especially between brainageR and DenseNet, and between PHOTON and Multimodal (r = 0.46–0.89). This suggests that models generally rank individuals similarly in terms of predicted age, even if they differ in scale or offset, which is captured by the ICC measure. It must be noted that correlation does not capture calibration differences; in other words, two models could be highly correlated in predicted age but differ in terms of correlation between brain-PADs. Any systematic offset in predicted age directly shifts the brain-PAD value, even if the rank order is preserved; brain-PAD is mean centered at 0, therefore the range of brain-PAD values have greater variance, so correlation is lower than when predicted ages are used.

*Agreement among algorithms as assessed by intraclass correlation coefficents*

Table 2.6. Intraclass Correlation Coefficients (ICCs) between predicted ages and corresponding 95% Confidence Interval

| Model 1 | Model 2 | ICC | N | 95% CI | p-value |
| --- | --- | --- | --- | --- | --- |
| Multimodal | PHOTON | 0.36 | 66 | 0.06-0.59 | < 0.01 |
| Multimodal | DenseNet | 0.17 | 66 | -0.09-0.44 | 0.137 |
| Multimodal | BrainageR | 0.43 | 66 | 0.05-0.71 | <0.05 |
| PHOTON | DenseNet | 0.78 | 117 | 0.21-0.91 | <0.01 |
| PHOTON | BrainageR | 0.73 | 117 | 0.63-0.80 | <0.0001 |
| DenseNet | BrainageR | 0.61 | 117 | 0.38-0.75 | <0.0001 |

Multimodal algorithm had lower agreement with the other models, particularly with DenseNet (ICC = 0.17), suggesting greater variability in predictions compared to the others. Note that given that Multimodal was trained on 51 HCs from our full sample of 73 HCs, we were only able to apply the model to the remaining 22 HCs to obtain a total of 66 predicted ages (HC = 22; BD = 44).

*Evaluation of Brain-PAD estimates as predictors of bipolar disorder*

Supplemental Table 2.4. Logistic regression model with brain-PADs across all three algorithms (HC = 73; BD = 44)

| Variable | Estimate | Std. Error | z value | p-value |
| --- | --- | --- | --- | --- |
| (Intercept) | –0.514 | 0.345 | –1.489 | 0.137 |
| PHOTON Brain-PAD | 0.064 | 0.026 | 2.429 | < 0.05 |
| BrainageR Brain-PAD | 0.005 | 0.033 | 0.144 | 0.886 |
| DenseNet Brain-PAD | –0.013 | 0.044 | –0.298 | 0.766 |

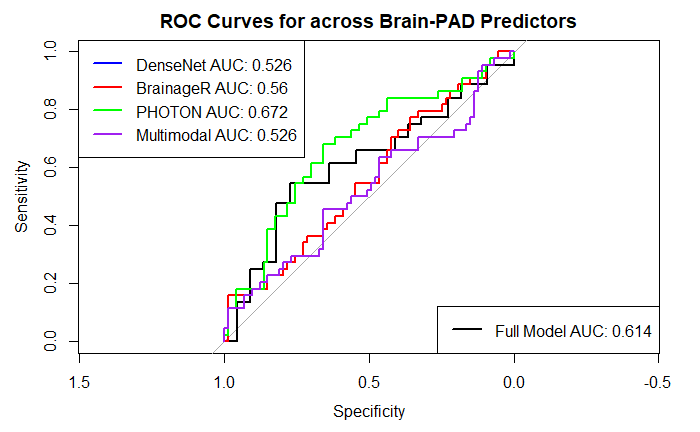

Supplemental Figure 2.4. Receiver Operating Characteristic (ROC) curves for distinguishing bipolar disorder (BD, N=44) from healthy controls (HC, N=22) based on raw Brain-PAD values derived from four brain age prediction algorithms: DenseNet, BrainageR, PHOTON and Multimodal.

Supplemental Table 2.5. Logistic regression model with brain-PADs across all four algorithms. (HC = 22; BD = 44)

| Variable | Estimate | Std. Error | z value | p-value |
| --- | --- | --- | --- | --- |
| (Intercept) | 0.738 | 0.603 | 1.224 | 0.221 |
| PHOTON Brain-PAD | 0.037 | 0.049 | 0.748 | 0.454 |
| BrainageR Brain-PAD | 0.029 | 0.052 | 0.571 | 0.568 |
| DenseNet Brain-PAD | –0.024 | 0.061 | –0.391 | 0.696 |
| Multimodal Brain-PAD | –0.028 | 0.044 | –0.651 | 0.515 |

When the Multimodal algorithm is included in the analyses (HC = 22; BD = 44), the linear regression model including brain-PAD estimates from PHOTON, BrainageR, DenseNet, and Multimodal showed no significant associations (AIC = 92.53). Overall, classification performance was poorer, with PHOTON again showing the highest AUC (0.672) compared to BrainAgeR (0.560), DenseNet (0.526), and Multimodal (0.526), and the full model yielding a lower AUC of 0.614. PHOTON Brain-PAD captures most of the signal available for discrimination in this dataset. The rank of algorithms based on this classification task is the same as when only three algorithms (Multimodal excluded) were evaluated.

*Relationship between chronological age and predicted age*

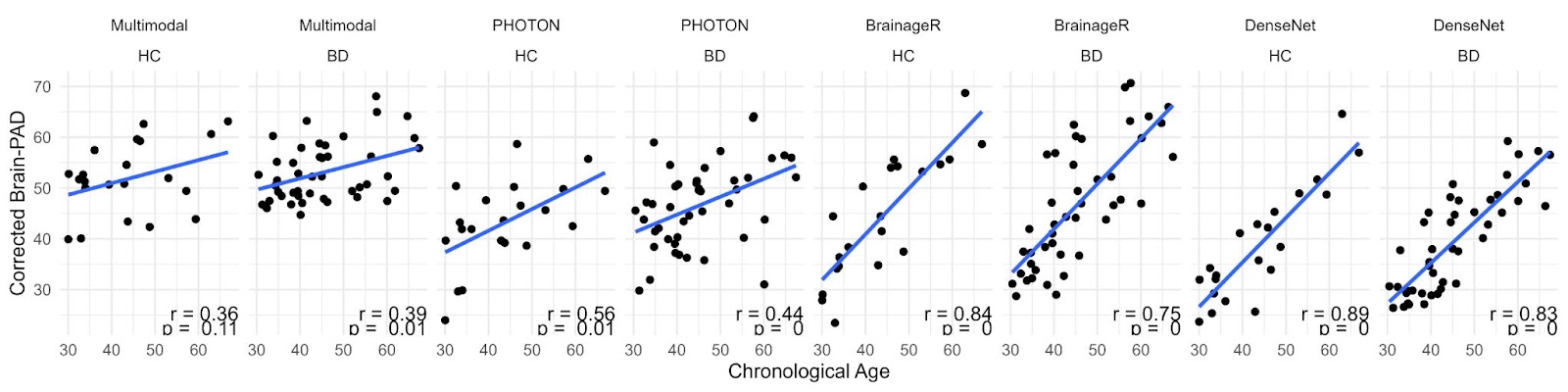

Supplemental Figure 2.5. Correlation between chronological age and predicted age across algorithms by group

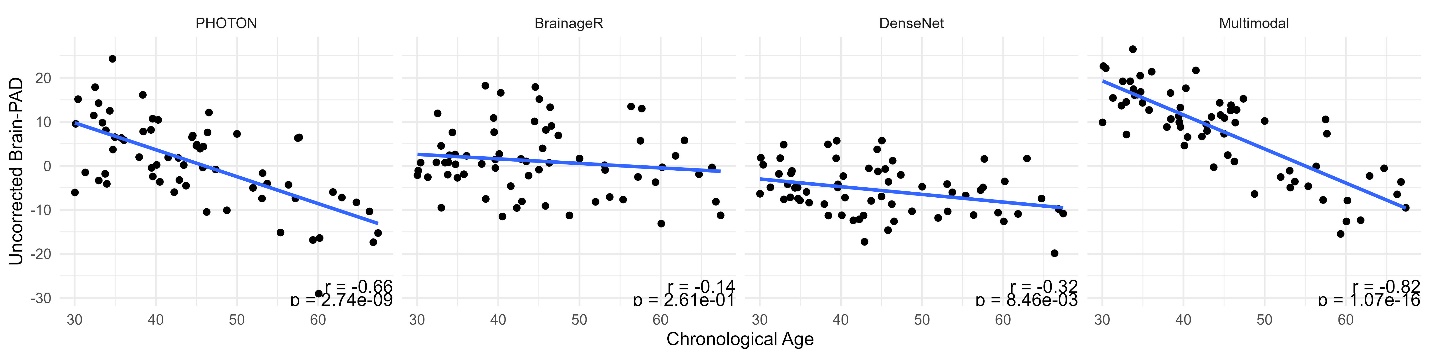

Supplemental Figure 2.6. Correlation between uncorrected brain-PAD and chronological age.

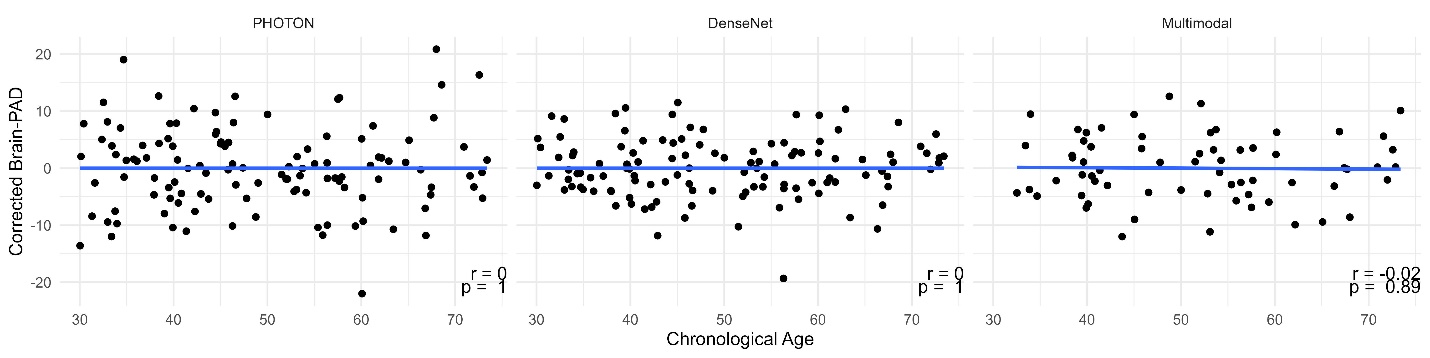

Supplemental Figure 2.7. Corrected Brain-PAD vs Chronological Age across the three algorithms

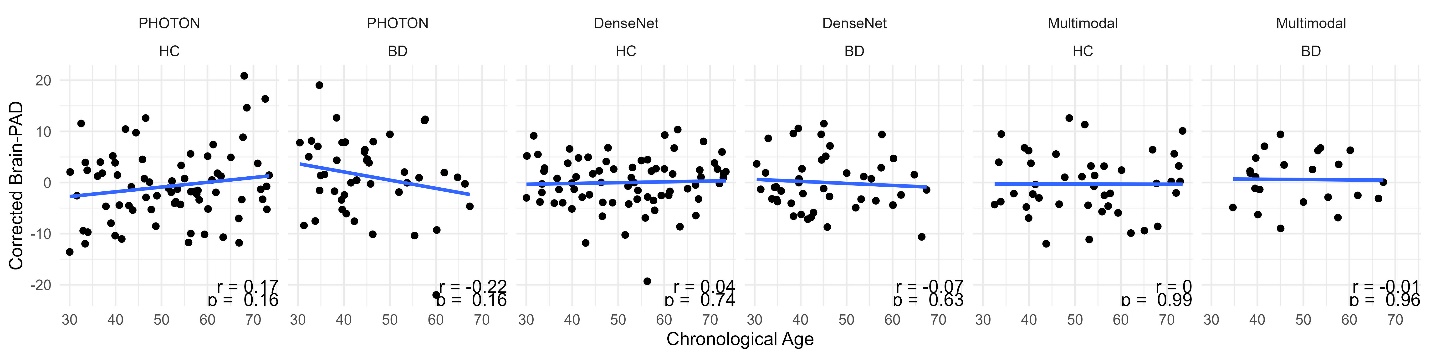

Supplemental Figure 2.8.  Corrected Brain-PAD vs Chronological Age by Diagnosis

DenseNet showed the strongest correlation between chronological age and predicted age followed by BrainageR, PHOTON and Multimodal. With the exception of the Multimodal algorithm, all algorithms demonstrated higher correlation among HC compared to BD. Prior to correction procedures, uncorrected Brain-PAD values showed strong negative correlations with chronological age across most algorithms (Multimodal: r = -0.82, PHOTON: r = -0.66, DenseNet: r = -0.32), with weaker and non-significant association in BrainageR (r = -0.14). These findings indicated that without correction, Brain-PAD is typically systematically biased by chronological age, with older individuals tending to show lower brain-PAD values (Supplemental Figure 2.6). To address this, we applied residualization procedures to correct for age-related bias for the algorithms that indicated bias. Following correction, we re-examined correlations between corrected Brain-PAD and chronological age. When examining the full sample without stratifying by diagnosis or sex, correlations between corrected Brain-PAD and chronological age were effectively zero across all algorithms (Multimodal: r = 0.07; PHOTON: r = 0.02; DenseNet: r ≈ 0), with non-significant p-values (Supplemental Figure 2.7). Across all algorithms, the correlations were substantially attenuated and close to zero (ranging from -0.14 to 0.17) within both the HC and BD groups. In HCs, the highest residual correlation was r = 0.17 for PHOTON, while BrainageR, DenseNet and Multimodal showed negligible correlations. Similar patterns were observed in BD participants, with very low or negative (-0.22 to -0.01) correlations (Supplemental Figure 2.8).

*Univariate correlation between input features of the multimodal algorithm and brain-PAD*

Structural changes such as cortical thinning and ventricular enlargement show the strongest associations with brain-PAD across HC and BD, followed by volumetric gray-matter and hippocampal measures. There is weak to no correlation between brain-PAD with functional (activation/CBF), surface-area and diffusion measures.

Supplemental Table 2.6. Correlation between brain-PAD and 87 input features in the multimodal algorithm.

|  | Feature | Correlation | P-Value |
| --- | --- | --- | --- |
| 1 | Left Hemisphere Total Average Cortical Thickness | -0.761 | < 0.001 |
| 2 | Left Frontal Lobe Average Cortical Thickness | -0.753 | < 0.001 |
| 3 | Right Hemisphere Total Average Cortical Thickness | -0.738 | < 0.001 |
| 4 | Right Lateral Parietal Lobe Cortical Thickness | -0.731 | < 0.001 |
| 5 | Right Parietal Lobe Average Cortical Thickness | -0.713 | < 0.001 |
| 6 | Left Superolateral Frontal Lobe Cortical Thickness | -0.712 | < 0.001 |
| 7 | Left Parietal Lobe Average Cortical Thickness | -0.7 | < 0.001 |
| 8 | Whole Brain Total Average Cortical Thickness | -0.695 | < 0.001 |
| 9 | Right Frontal Lobe Average Cortical Thickness | -0.683 | < 0.001 |
| 10 | Right Superolateral Frontal Lobe Cortical Thickness | -0.676 | < 0.001 |
| 11 | Right Medial Parietal Lobe Cortical Thickness | -0.672 | < 0.001 |
| 12 | Left Lateral Parietal Lobe Cortical Thickness | -0.667 | < 0.001 |
| 13 | Left Medial Parietal Lobe Cortical Thickness | -0.64 | < 0.001 |
| 14 | Right Medial Inferior Frontal Lobe Cortical Thickness | -0.584 | < 0.001 |
| 15 | Left Medial Inferior Frontal Lobe Cortical Thickness | -0.502 | < 0.001 |
| 16 | Right Lateral Ventricle Volume | 0.495 | < 0.001 |
| 17 | Left Lateral Ventricle Volume | 0.471 | < 0.001 |
| 18 | Right Hippocampal & Parahippocampal Volume | -0.466 | < 0.001 |
| 19 | Right Inferior Lateral Ventricle Volume | 0.415 | < 0.001 |
| 20 | Right Hemisphere Total Gray Matter Volume | -0.414 | < 0.001 |
| 21 | Right Hippocampus Volume | -0.407 | < 0.001 |
| 22 | Whole Brain Total Gray Matter Volume | -0.399 | < 0.001 |
| 23 | Right Parahippocampal Volume | -0.397 | < 0.001 |
| 24 | Left Hemisphere Total Gray Matter Volume | -0.395 | < 0.01 |

Supplemental Table 2.6 (Continued)

| 25 | Left Inferior Lateral Ventricle Volume | 0.395 | < 0.01 |
| --- | --- | --- | --- |
| 26 | Left Hippocampus Volume | -0.37 | < 0.01 |
| 27 | Left Hippocampal & Parahippocampal Volume | -0.355 | < 0.01 |
| 28 | Right Anterior Cingulate Deactivation (Task Contrast 2v1) | 0.351 | < 0.01 |
| 29 | Right Middle Frontal Gyrus Activation (Task Contrast 2v1) | -0.328 | < 0.01 |
| 30 | Left Periventricular Hyperintensity Volume (FLAIR, mm³) | 0.321 | < 0.01 |
| 31 | Right Parietal Lobule Activation (Task Contrast 2v1) | -0.309 | < 0.05 |
| 32 | Left Middle Frontal Gyrus Activation (Task Contrast 2v1) | -0.295 | < 0.05 |
| 33 | Corpus Callosum Splenium FA (TBSS) | -0.262 | < 0.05 |
| 34 | Corpus Callosum Genu FA (TBSS) | -0.252 | < 0.05 |
| 35 | Right Medial Inferior Frontal Lobe Surface Area | 0.238 | NS |
| 36 | Right Superior Longitudinal Fasciculus FA (TBSS) | -0.224 | NS |
| 37 | Right Periventricular Hyperintensity Volume (FLAIR, mm³) | 0.223 | NS |
| 38 | Left Medial Inferior Frontal Lobe Surface Area | 0.214 | NS |
| 39 | Left Parahippocampal Volume | -0.214 | NS |
| 40 | Right Cingulum–Hippocampus FA (JHU Atlas) | -0.197 | NS |
| 41 | Left Superior Longitudinal Fasciculus FA (TBSS) | -0.188 | NS |
| 42 | Left Deep White Matter Hyperintensity Volume (FLAIR, mm³) | 0.163 | NS |
| 43 | Left Parietal Lobe CBF | 0.162 | NS |
| 44 | Left Lateral Parietal Lobe CBF | 0.16 | NS |
| 45 | Left Supramarginal Gyrus (Parietal) DMS Activation | 0.16 | NS |
| 46 | Right Uncinate Fasciculus FA (TBSS) | -0.16 | NS |
| 47 | Left Frontal Lobe Surface Area | 0.155 | NS |
| 48 | Left Uncinate Fasciculus FA (TBSS) | -0.152 | NS |
| 49 | Right Medial Parietal Lobe CBF | 0.14 | NS |
| 50 | Left Hippocampus CBF (Mean) | 0.132 | NS |
| 51 | Left Medial Inferior Frontal Lobe CBF | -0.124 | NS |
| 52 | Right Superolateral Frontal Lobe Surface Area | 0.122 | NS |

Supplemental Table 2.6 (Continued)

| 53 | Left Hemisphere Total Surface Area | 0.12 | NS |
| --- | --- | --- | --- |
| 54 | Corpus Callosum Body FA (TBSS) | -0.107 | NS |
| 55 | Right Hemisphere CBF | 0.105 | NS |
| 56 | Right Deep White Matter Hyperintensity Volume (FLAIR, mm³) | 0.097 | NS |
| 57 | Left Superolateral Frontal Lobe Surface Area | 0.095 | NS |
| 58 | Right Lateral Parietal Lobe CBF | 0.093 | NS |
| 59 | Right Frontal Lobe CBF | 0.089 | NS |
| 60 | Right Medial Inferior Frontal Lobe CBF | 0.083 | NS |
| 61 | Right Frontal Lobe Surface Area | 0.081 | NS |
| 62 | Left Medial Parietal Lobe Surface Area | 0.08 | NS |
| 63 | Whole Brain Total Surface Area | 0.077 | NS |
| 64 | Left Middle Frontal Gyrus DMS Activation | -0.068 | NS |
| 65 | Left Lateral Parietal Lobe Surface Area | -0.063 | NS |
| 66 | Right Middle Frontal Gyrus DMS Activation | -0.055 | NS |
| 67 | Task Positive Network Z Correlation | -0.054 | NS |
| 68 | Right Supramarginal Gyrus (Parietal) DMS Activation | 0.046 | NS |
| 69 | Left Superiolateral Frontal Lobe CBF | 0.044 | NS |
| 70 | Right Parietal Lobe CBF | 0.039 | NS |
| 71 | Right Parahippocampal CBF (Mean) | 0.035 | NS |
| 72 | Right Hippocampus CBF (Mean) | 0.035 | NS |
| 73 | Right Superolateral Frontal Lobe CBF | 0.033 | NS |
| 74 | Left Frontal Lobe CBF | 0.031 | NS |
| 75 | Left Parietal Lobe Surface Area | -0.026 | NS |
| 76 | Right Medial Parietal Lobe Surface Area | 0.025 | NS |
| 77 | DMN Resting-State Z Correlation | -0.024 | NS |
| 78 | Left Medial Parietal Lobe CBF | -0.021 | NS |
| 79 | Right Parietal Lobe Surface Area | 0.021 | NS |
| 80 | Right Hippocampal & Parahippocampal Volume | -0.019 | NS |

Supplemental Table 2.6 (Continued)

| 81 | Right Hemisphere Total Surface Area | -0.018 | NS |
| --- | --- | --- | --- |
| 82 | Left Cingulum–Hippocampus FA (JHU Atlas) | 0.012 | NS |
| 83 | Left Hippocampal & Parahippocampal Volume | 0.008 | NS |
| 84 | Left Parahippocampal CBF (Mean) | -0.004 | NS |
| 85 | Right Lateral Parietal Lobe Surface Area | 0.003 | NS |
| 86 | Left Hemisphere CBF | 0.001 | NS |
| 87 | Whole Brain Gray Matter CBF (nz Mean) | 0.000 | NS |

Supplemental Table 2.7

| Modality | Features |
| --- | --- |
| FLAIR - White Matter Hyperintensity | Left Periventricular Hyperintensity Volume, Right Periventricular Hyperintensity Volume, Left Deep White Matter Hyperintensity Volume, Right Deep White Matter Hyperintensity Volume |
| White Matter FA (JHU Atlas) | Right Cingulum–Hippocampus FA, Left Cingulum–Hippocampus FA |
| White Matter FA (TBSS) | Corpus Callosum Splenium FA, Corpus Callosum Genu FA, Corpus Callosum Body FA, Right Superior Longitudinal Fasciculus FA, Left Superior Longitudinal Fasciculus FA, Right Uncinate Fasciculus FA, Left Uncinate Fasciculus FA |
| Gray Matter CBF | Left Parietal Lobe CBF, Left Lateral Parietal Lobe CBF, Right Medial Parietal Lobe CBF, Left Hippocampus CBF (Mean), Left Medial Inferior Frontal Lobe CBF, Right Hemisphere CBF, Right Lateral Parietal Lobe CBF, Right Frontal Lobe CBF, Right Medial Inferior Frontal Lobe CBF, Left Superiolateral Frontal Lobe CBF, Right Parietal Lobe CBF, Right Parahippocampal CBF (Mean), Right Hippocampus CBF (Mean), Right Superolateral Frontal Lobe CBF, Left Frontal Lobe CBF, Left Medial Parietal Lobe CBF, Left Hemisphere CBF, Left Parahippocampal CBF (Mean), Whole Brain Gray Matter CBF (nz Mean) |
| Gray Matter Average Thickness | Left Hemisphere Total Average Cortical Thickness, Left Frontal Lobe Average Cortical Thickness, Right Hemisphere Total Average Cortical Thickness, Right Lateral Parietal Lobe Cortical Thickness, Right Parietal Lobe Average Cortical Thickness, Left Superolateral Frontal Lobe Cortical Thickness, Left Parietal Lobe Average Cortical Thickness, Whole Brain Total Average Cortical Thickness, Right Frontal Lobe Average Cortical Thickness, Right Superolateral Frontal Lobe Cortical Thickness, Right Medial Parietal Lobe Cortical Thickness, Left Lateral Parietal Lobe Cortical Thickness, Left Medial Parietal Lobe Cortical Thickness, Right Medial Inferior Frontal Lobe Cortical Thickness, Left Medial Inferior Frontal Lobe Cortical Thickness |
| Gray Matter Surface Area | Right Medial Inferior Frontal Lobe Surface Area, Left Medial Inferior Frontal Lobe Surface Area, Left Frontal Lobe Surface Area, Right Superolateral Frontal Lobe Surface Area, Left Hemisphere Total Surface Area, Left Superolateral Frontal Lobe Surface Area, Left Medial Parietal Lobe Surface Area, Whole Brain Total Surface Area, Left Lateral Parietal Lobe Surface Area, Right Middle Frontal Gyrus Surface Area, Right Parietal Lobe Surface Area, Right Lateral Parietal Lobe Surface Area, Right Hemisphere Total Surface Area |
| Gray Matter Volume | Right Hippocampal & Parahippocampal Volume, Right Hemisphere Total Gray Matter Volume, Right Hippocampus Volume, Whole Brain Total Gray Matter Volume, Right Parahippocampal Volume, Left Hemisphere Total Gray Matter Volume, Left Hippocampus Volume, Left Hippocampal & Parahippocampal Volume, Left Parahippocampal Volume |
| Resting State Functional Connectivity | Task Positive Network Z Correlation, DMN Resting-State Z Correlation |
| Task-Based fMRI (N-back / DMS) | Right Anterior Cingulate Deactivation (Task Contrast 2v1), Right Middle Frontal Gyrus Activation (Task Contrast 2v1), Right Parietal Lobule Activation (Task Contrast 2v1), Left Middle Frontal Gyrus Activation (Task Contrast 2v1), Left Supramarginal Gyrus (Parietal) DMS Activation, Left Middle Frontal Gyrus DMS Activation, Right Middle Frontal Gyrus DMS Activation, Right Supramarginal Gyrus (Parietal) DMS Activation |

*Multimodal algorithm development, validation and evaluation*

BD is characterized by interconnected structural and functional brain abnormalities. Widespread WM microstructural alterations identified via DTI suggest disruption of key WM tracts, which likely underlies both altered functional connectivity patterns and CBF changes. Concurrently, meta-analyses of T2-weighted imaging reveal an increased prevalence of WM hyperintensities, indicating small-vessel pathology that may further impair WM integrity. Functional MRI studies show aberrant connectivity within fronto-limbic and large-scale networks cortical thinning and subcortical volume reductions gray matter (GM) alterations. As a result, despite the leave-out-one-modality analysis suggesting that the model trained without cortical thickness features performed the best, we opted to include cortical thickness features in model training (Supplementary Table 2.8). After all, cortical thinning is a hallmark of normal brain aging and a sensitive structural biomarker of cognitive decline in aging and BD (Abe et al., 2022, Hibar et al., 2018). By developing the Multimodal algorithm, we focus on examining whether doing so improves sensitivity to subtle local cohort-specific effects and reduces domain shift relative to off-the-shelf algorithms. For instance, differences in scanner (e.g., field strength, echo times), pre-processing methods (e.g., skull stripping, motion correction), demographics (e.g., age range, sex distribution) and strictness of inclusion criteria (e.g., definition of radiologically ‘normal’) contribute to domain-shift effects that decrease the accuracy of brain-PAD estimates.
